# A Bottom-Up Approach to Fungal Plasma Membrane Model: Lipid Mixture Design and Biophysical-Mechanical Characterization

**DOI:** 10.64898/2026.08.08.743690

**Authors:** Maxime Kucharski, Zuzanna Kubicka, Dominik Drabik

## Abstract

The rising incidence of invasive fungal diseases emphasizes the need for novel therapeutic strategies, including membrane-targeting antifungal agents, which require representative lipid models for detailed molecular-level studies. In this work, we propose a consensus quinary fungal plasma membrane model based on lipidomic literature data, specifically PC:PE:PI:PA:PS phospholipid model with ratio of 44:29:13:8:6. Using a bottom-up approach, we characterized the biophysical properties of this system – with particular emphasis on mechanical parameters such as bending rigidity and area compressibility – by combining molecular dynamics simulations with experimental flicker-noise and ATR-FTIR spectroscopies. Furthermore, we investigated the effect of two key non-phospholipid components: ergosterol and triacylglycerols. Biophysical analysis revealed that DPPI and its specific interactions with DSPS induced the most substantial deviations in baseline membrane parameters, particularly area per lipid, membrane thickness, and area compressibility, while DSPS influenced bending rigidity change and DLiPA primarily affected lipid packing defects. In addition, ergosterol and TGs were found to influence all of the investigated parameters to different degree. Notably, the overall biophysical profile of the proposed FPMM closely mimicked that of natural vesicles derived from yeast lipid extracts, establishing this model may provide a reliable platform for studying fungal membrane biophysics and lipid-targeting interactions.

## 1. Introduction

Invasive fungal diseases are on the rise, particularly among immunocompromised individuals. Limited diagnostic tools, treatments, and the emergence of antifungal resistance hinder effective management of these infections. They impose a significant public health burden, with mortality rates ranging from 30% to 90% depending on the specific pathogen and patient population (1). Fungal infections receive little attention and resources, resulting in a lack of quality data on fungal disease distribution and antifungal resistance patterns. Yet, the problem of antimicrobial resistance is primarily discussed in the context of bacteria, despite the increasing threat of fungal infections. Fortunately, in 2022, the World Health Organization (WHO) published its first-ever list of fungal priority pathogens, identifying 19 fungi that pose the greatest threat to public health (2). Climate change may further exacerbate the issue, as rising temperatures could facilitate the evolution and emergence of new fungal pathogens. This emphasizes the need to develop new antifungal substances. A way to address this is to see how this is handled in the case of antimicrobial substances. One promising strategy is the targeting of bacterial membranes. This approach has proven successful with such substances as octenidine (3) and general class of cationic gemini molecules (4). A similar strategy could be applied to fungi, but requires a suitable lipid membrane model. While fungal mycelium are well-suited for physiological and biochemical studies, fungal membranes remain relatively understudied (5). Generalizations about fungal membrane lipid composition are often based on yeast cells and a limited number of filamentous fungi. Typically, a fungal cell plasma membrane comprises phospholipids and ergosterol. The cell wall, composed of chitin, *β*-glucan, and mannoproteins layers, provides rigidity and protection The plasma membrane of fungi is a key target for understanding and combating fungal infections. Lipids, especially glycolipids and ergosterol, play crucial roles in fungal virulence, drug resistance, and adaptation to environmental stress (6). Fungal membranes differ significantly from bacterial membranes in their lipid composition. Phosphatidylcholines (PC) and phosphatidylethanolamines (PE) are the primary phospholipids in fungi, while ergosterol is a predominant sterol component (7; 8; 9; 10). The relative abundance of these lipids can vary depending on the fungal species and environmental conditions. The remaining 20% belongs to anionic phospholipids: phosphatidylserines and phosphatidylinositols (11). The varying lipid composition of fungal membranes influences their properties and functions. Recent studies have highlighted the significance of the physical properties of membrane lipids in pathogenic fungi (6). Lipid microdomains, enriched in glycosphingolipids and sterols, can concentrate virulence factors and contribute to pathogenicity (12; 13; 14; 15). Changes in membrane fluidity, resulting from changes in lipid homeostasis, have been shown to affect drug resistance in Candida species (16; 17). Thus, the physical properties of the plasma membrane appear to influence the outcome of fungal infections. This manuscript aims to propose consensual lipid model mixture representing the plasma membranes of several fungal species, with a specific focus on the distribution of ergosterol and triacylglycerols. Utilizing available lipidomic data, we developed a representative fungal membrane model composed of five distinct lipid species. This system was subsequently characterized biophysically through a complementary approach combining molecular dynamics (MD) simulations with experimental techniques, namely, flicker-noise spectroscopy and attenuated total reflection Fourier-transform infrared (ATR-FTIR) spectroscopy. Furthermore, given the biological relevance of ergosterol and triacylglycerols in fungal physiology, we investigated the individual effects of these components on membrane properties.

## 2. Results and Discussion

### 2.1. Bottom-up fungal membrane model

To design an accurate and representative fungal plasma membrane model (FPMM), an analysis of available lipidomic data is necessary. A review of this literature data is summarized in Table 1. Based on these reported values, the average compositions for each lipid species were calculated, yielding a baseline phospholipid ratio of PC:PE:PI:PA:PS (44:29:13:8:6); notably, both lyso-PC and lyso-PE were omitted from the model due to their negligible fractions. However, selecting the lipid headgroups addresses only one part of model development; the second aspect is the acyl chain composition. Given that fungal plasma membranes typically exhibit phospholipid acyl chains ranging from 16 to 18 carbons (23), with varying degrees of unsaturation, our selection directly reflects these physiological constraints. To capture this diversity, we utilized specific phospholipids varying in both chain length and saturation: DPPI (16:0), DYPE (16:1), POPC (16:0–18:1), DLiPA (18:2), and DSPS (18:0). To investigate the biophysical contribution of each lipid species to the membrane model, we adopted a bottom-up approach. Specifically, we initiated both experimental and simulation approaches with a simplified two-component POPC:DYPE membrane, which was then incrementally expanded to generate increasingly complex ternary, quaternary, and quinary systems constant molar ratio of individual lipids. We acknowledge that the use of lipid species with distinct acyl-chain compositions introduces a potential confounding factor, making it difficult to unambiguously attribute observed changes solely to given lipid aspect. Nevertheless, this approach was adopted to balance the complexity of the lipid model with the objective of capturing key features of biologically relevant membrane compositions while limiting the number of lipid components.

**Table 1:** Lipidomic data of available fungi membrane. Apart from *istoplasma capsulatum*, they all follow similar distribution. Membranes of *Histoplasma* and *Penicillium* additionally consisteded of 45% and 30% triacylglycerol, respectively.

| Fungi Species | lysoPC | PC | lysoPE | PE | PI | PS | PA | PG | Source |
| --- | --- | --- | --- | --- | --- | --- | --- | --- | --- |
| *Candida auris*<br>(drug susceptible)<br>[%] (nmol/mg) | 5.5<br>(13.65) | 44<br>(109.33) | 2.35<br>(5.84) | 21.5<br>(53.52) | 12.9<br>(32.09) | 2.9<br>(7.25) | 9.15<br>(22.73) | 1.65<br>(4.11) | (18) |
| *Candida auris*<br>(drug resistant)<br>[%] (nmol/mg) | 3.3<br>(15.52) | 40<br>(186.8) | 1.9<br>(8.75) | 32.8<br>(153.2) | 11.6<br>(54.4) | 2.6<br>(12.0) | 5.45<br>(5.45) | 2.47<br>(11.53) | (18) |
| *Cryptococcus*<br>[%] | no data | 48 | 0.6 | 24.4 | 7.9 | 12.6 | 2 | 0.2 | (19) |
| *Saccharomyces cerevisiae*<br>[%] | no data | 40 | no data | 25 | 20 | 5 | 0 | 0 | (20) |
| *Penicillium digitatum*<br>[%] | 2.6 | 30.5 | 2.2 | 3.5 | 4.35 | 13.9 | 10.5 | 1.2 | (21) |
| *Histoplasma capsulatum*<br>[%] | no data | 5 | no data | 19 | 11 | 3 | 8 | 4 | (20) |
| *Neurospora crassa*,<br>plasma membrane<br>[%] | no data | 46.43 | no data | 39.38 | 15.18 | 1 | no data | no data | (22; 5) |

Abbreviations: lysoPC: 1-lysophosphatidylcholine, PC: phosphatidylcholine, lysoPE: 2-linoleoyl-sn-glycero-3- phosphoethanolamine, PE: phosphoethanolamine, PI: phosphatidylinositol, PS: phosphatidylserine, PA: phosphatidic acid, PG: phosphatidylglycerol.

Interestingly, most of the membrane systems’ properties remained constant during the course of model bottom-up building. There is significant difference in MT in between FPMM and base mixture (4.01 ± 0.01 vs 3.88 ± 0.01), as shown in Fig 1A. When looking on the data from tertiary system, neither DSPS nor DLiPA affected MT, however DPPI increased the MT significantly, but not the the level of quinary FPMM. Interestingly, in quaternary system when both DPPI and DSPS lipids are present, the MT is higher than for the tetriary system with DPPI alone, indicating the interaction between those two lipid species. Interestingly, system with DSPS and DLiPA had higher MT than the respective tetriary systems, showing that either DSPS presence or PS/PA interactions resulted in increase of MT. However, this effect was not present in tetriary system with DSPS, which indicates that presence of DLiPA is necessary. The same goes for area per lipid (APL) - significant difference in APL in between FPMM and base mixture (59.6 ± 0.2 vs 57.9 ± 0.3). Although the simultaneous increase of MT and decrease of APL are common signatures of a gel-like transition, the underlying lateral domain organization of this system is addressed below. Furthermore, interdigitation remained constant throughout most of the system (with small decrease for systems with DPPI), so it is unlikely that this is due to interactions between the leaflets. To this end, we hypothesize that interactions between saturated and unsaturated lipids are a primary driver of these changes (24; 25).

**Figure 1:**
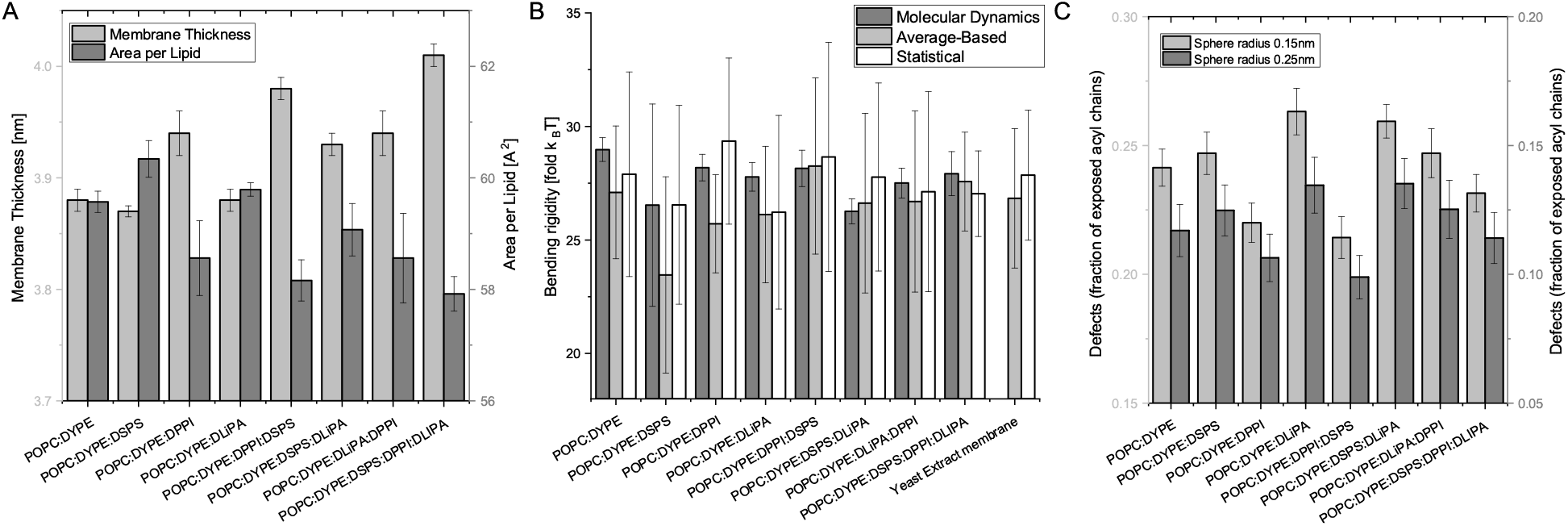
Biophysical characteristics of the investigated lipid membrane systems. (A) Membrane thickness (MT) and area per lipid (APL) determined from molecular dynamics (MD) simulations. (B) Bending rigidity (κ) determined from MD simulations and experimentally via flicker-noise spectroscopy using two different calculation approaches. (C) Acyl chain accessibility sphere analysis used to quantify membrane packing defects for the investigated systems at two probe radii (*r* = 0.15 nm and *r* = 0.25 nm).

These structural changes were independently confirmed via ATR-FTIR spectroscopy. Comparing the spectra of final mixtures to individual lipid standards confirmed the presence of all component lipids, while observed band shifts pointed to specific intermolecular interactions (26). In the PC:PE:PS system, the absence of the characteristic PS band at 1620 cm^−1^ suggests interactions affecting the serine carboxylate group. In the PC:PE:PI mixture, the 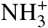 doublet shift of the PE head-group was more pronounced than in PC:PE:PS, indicating a greater perturbation of the PE headgroup environment by PI, potentially reflecting stronger hydrogen-bonding and/or electrostatic interactions. On the contrary, no distinct 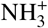 doublet was observed in the 1580–1680 cm^−1^ region for PC:PE:PA. Furthermore, the PA-related at 935 cm^−1^ was absent in both PC:PE:PA and PC:PE:PA:PI, yet re-emerged in PC:PE:PS:PA and PC:PE:PI:PA:PS, suggesting that PS modifies the local PA headgroup environment. Given the comparable PA order parameters, this effect is unlikely to arise from changes in acylchain ordering and may instead reflect altered electrostatic interactions or PA protonation (27). Correlating the spectroscopic data with physical parameters, the ratio of asymmetric to symmetric CH_2_ stretching vibrations (∼2920 cm^−1^ and ∼2850 cm^−1^, respectively) serves as an indicator of *gauche* defects, where higher ratios reflect increased fluidity, larger APL, and reduced MT. This ratio correlated well with APL changes across most systems, though the structural alterations were overestimated for PC:PE:PA and PC:PE:PI:PS. In PC:PE:PA, this discrepancy likely due to effect of PA on the PE headgroup, whereas in PC:PE:PI:PS, most likely electrostatic repulsion between the negatively charged PI and PS headgroups forces additional lateral spacing. Finally, observed MT changes correlated more strongly with isolated shifts in the 2920 cm^−1^ band alone. Spectra are presented in Fig S1.

The investigated membrane systems exhibited high baseline mechanical stability, although distinct compositional deviations were observed. The results for bending rigidity (*κ*), which quantifies the energetic cost of membrane deformation, were obtained from both molecular dynamics simulations and experimental fluctuation analysis, as presented in Fig 1B. Based on the data from the ternary systems, the presence of DSPS decreased the bending rigidity, whereas the incorporation of either DLiPA or DPPI resulted in an increased rigidity. Interestingly, in the quaternary system containing both DSPS and DLiPA, the bending rigidity remained comparable to that of the ternary system with DSPS alone; the stiffening effect imparted by DLiPA in the ternary mixture was effectively abolished. Such effect was not observed in the quaternary system combining DSPS and DPPI. This mechanical softening is most likely driven by specific PA and PS headgroup interactions. Phosphatidic acid (PA) possesses a unique capacity to participate in intermolecular hydrogen bonding with neighboring lipid headgroups. These localized hydrogen-bonding networks, alongside the resulting alterations in the interfacial hydration layer, significantly modulate the lateral pressure profile across the bilayer. Consequently, the membrane compensates for these altered lateral stresses by becoming structurally softened and more amenable to bending (28). Furthermore, the bending rigidity of the membrane derived from the native yeast polar extract was in agreement with our proposed fungal membrane model. This closely matched behavior confirms that, at least from a mechanical standpoint, our proposed membrane model follows the physical behavior of native yeast plasma membranes.

Second mechanical property, the area compressibility modulus (*K*_*A*_), which measures the membrane’s resistance to lateral expansion or compression, remained relatively constant across most investigated systems. A significant increase in *K*_*A*_ was detected exclusively in systems containing DPPI, which aligns with the known rigidifying effects of this lipid species. However, given the acyl chain lenght, is is also possible that this effect results from acyl chain lenght mismatch. Because this elevated *K*_*A*_ was unique to DPPI-enriched bilayers, it suggests that this specific lipid in the investigated systems is primarily responsible for modulating lateral membrane resilience, with no observable cooperative interactions from other lipids affecting this property. Additionally, dynamic parameters such as lateral diffusion coefficients were largely unaffected by changes in composition, exhibiting only a minor decrease upon the incorporation of DPPI. Numerical values for all evaluated parameters are provided in Table S1.

The mechanical characterization was further complemented by ATR-FTIR spectroscopic analysis. Spectral shifts in the 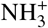 region of PE (∼1673 cm^−1^) signify variations in inter-headgroup hydrogen bonding. Robust headgroup hydrogen-bonding networks stiffen the interfacial plane, thereby increasing membrane bending rigidity. This relationship held true for most compositions, correlating well across the dataset with the exception of the PC:PE:PS system. Interestingly, *K*_*A*_ exhibited no correlation with either the symmetric CH_2_ stretching vibrations (*ν*_*s*_(CH_2_)) or the 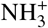 PE headgroup region. Instead, a correlation was identified for the ester carbonyl (C=O) stretching vibrations originating from the glycerol backbone region (∼1730 cm^−1^). Nevertheless, both mechanical responses were clearly reflected in the spectroscopic data.

The characteristics of system wouldn’t be completed without distribution of lipid packing defects analysis. It is important, yet less straightforwardly established, membrane property. To quantify this property, we estimated the acyl chain-accessible surface regions following the approach proposed by Boyd et al. (29). A map of accessible regions was generated for each frame, and spheres of a predefined radius were fitted into the map to prevent overestimating defect sizes caused by local thermal fluctuations of lipids. Results can be seen in Fig 1C. Based on these calculations, incorporation of the DLiPA lipid led to an increase in packing defects. This is expected, as its smaller headgroup leaves the underlying acyl chain region more exposed (30). On contrary, the fully saturated DPPI lipid reduced packing defects by inducing tighter membrane packing. This is likewise consistent with the expectation that at such temperature, the saturated acyl chains remain highly ordered, allowing adjacent lipids to pack more closely. Finally, DSPS did increase acyl chain accesibility, but it was not significantly different from referenced system; this trend persisted in quaternary systems containing DLiPA or DPPI, where acyl chain exposure closely mirrored that of the corresponding ternary systems. Interestingly, the quaternary system containing both DSPS and DPPI exhibited significantly lower acyl-chain exposure than would be expected from a simple additive ratio of the respective ternary systems. This suggests that the presence of DSPS may further modify membrane organization, potentially due to the acyl-chain length mismatch between DPPI and DSPS. Both quaternary systems with DLiPA, exhibited higher acyl chain exposure than calculated from simple additive. This indicates that PA exerts a pronounced influence on lipid packing defects that is not counteracted by the presence of DPPI. This non-additive behavior is most likely driven by specific intermolecular interactions of PA with either PI, and PS headgroups, or both. This mechanism is supported by analyzing the local lipid environment within a 12 Å cutoff of PA molecules in FPMM. A two-fold increase in the fraction of PA lipids in close proximity to both PI and PS was observed between the initial state and the final trajectory replicas (increasing from 1.7% to 3.7%). Concurrently, the fraction of PA in the vicinity of PS alone (without PI) increased from 4.3% to 6.8%. In contrast, the fraction of PA neighboring PI alone (without PS) decreased from 16.3% to 12.4%, pointing to repulsive interactions between these negatively charged lipid headgroup types (see Fig S2A).

One intriguing aspect is domain occurrence within the proposed membrane model. In biological systems, fungal plasma membranes frequently exhibit lipid segregation (31; 32). Across all investigated systems, two distinct features were observed. First, in the tertiary system containing DPPI (as well as in tetriary system containing DPPI and DSPS), local buds with increased fluorescence intensity were detected (Fig 2A). Second, when DPPI was present in a tertiary system alongside DLiPA, distinct fluorescence-free regions were visible (Fig 2B). This was observed with both of investigated probes. Similar domains were observed both in the final quinary fungal membrane model and in vesicles prepared from total fungal lipid extracts (Fig 2C). Interestingly, no lipid species exhibited an elevated lipid order parameter (see Fig S3), rendering the possible explanation of liquid-ordered domain formation unlikely. Instead, the observed domain segregation is most likely driven by hydrogen-bonding networks between PE and PI lipids resulting in local buds (33), which subsequently evolve into fluorescence-free regions upon the addition of PA, likely mediated by interactions among PE, PI, and PA lipids. Indeed, analysis of hydrogen bonding within the system reveals a substantial proportion of hydrogen bonds forming between DPPI and DYPE residues. In the PC:PE:PI system, this interaction represented the highest observed fraction at 43.9%, which is high given the relative lipid abundance. By comparison, in the PC:PE:PA system, hydrogen bonds between DLiPA and DYPE comprised only 28.6% of total inter-lipid hydrogen bonds. In the 4- and 5-component systems, the relative contributions for DPPI–DYPE and DLiPA–DYPE pairs remained consistent at approximately 37% and 15%, respectively. Direct hydrogen bonding between DLiPA and DPPI was minor, occurring in only 6% of cases (Fig S2B). Together, these findings suggest that specific hydrogen-bonding networks contribute to the observed domain segregation.

**Figure 2:**
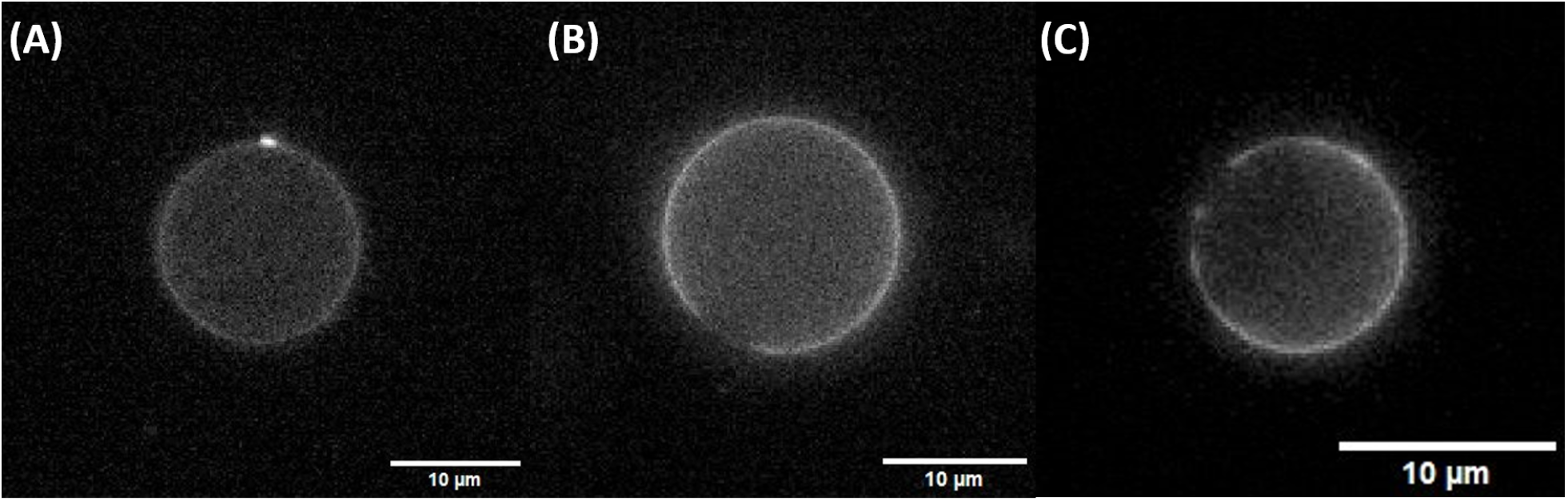
Domain segregation observed in the investigated lipid vesicles. (A) Local buds with increased fluorescence intensity detected in the POPC:DYPE:DPPI:DSPS system. (B) Distinct fluorescence-free regions observed in the POPC:DYPE:DPPI:DLiPA system. (C) Similar fluorescence-free regions observed in vesicles formed from yeast polar lipid extracts.

Collectively, these findings indicate that proposed POPC:DYPE:DSPS:DLiPA:DPPI lipid mixture can be used to accurately model biophysical characteristic of FPMM. Furthemore, among all investigated lipids, the independent presence of DPPI - or its specific interaction with DSPS - induced the most substantial deviations in baseline membrane properties, particularly regarding APL, MT and *K*_*A*_. Conversely, DSPS played a more dominant role specifically in modulating membrane bending rigidity. In contrast, the highly unsaturated DLiPA did not significantly alter the baseline physical characteristics of the membrane, though had the most significant impact on membrane defects.

### 2.2. Effect of Ergosterol

Sterols are essential components of eukaryotic plasma membranes, playing a critical role in regulating membrane integrity, fluidity, and permeability, while frequently partitioning into specific domains. Ergosterol is widely recognized as the primary fungal sterol, often serving as a biomarker for fungi in medical and environmental analyses (10; 9). The share of ergosterol in the lipid pool of the fungal membrane varies substantially depending on the species, developmental stage, and environmental conditions, ranging from near 0% to over 50% of all membrane lipids. Adjustments in plasma membrane ergosterol content contribute significantly to fungal adaptation under diverse environmental stresses, including elevated temperatures, osmotic shock, and oxidative stress (34). The biophysical effect of ergosterol is consistent across most investigated parameters and similar to the behavior of other eukaryotic sterols such as cholesterol. The sole exception is MT, which increases initially but subsequently decreases between 30% and 50% ergosterol concentration (from 4.12 ± 0.01 nm to 4.06 ± 0.01 nm) (Fig 3A). In contrast, a decrease in APL alongside an increase in both mechanical properties — bending rigidity and *K*_*A*_ — was observed, which is a well-established condensing and rigidifying effect of sterols in lipid bilayers (35). The exact properties are also reflected in existing data regarding ergosterol influence on lipid membranes (36; 37; 38). Notably, the results are in agreement with previous MD simulations concerning similar membrane models to presented quinary system, where reduction of APL accompanied by increase in *K*_*A*_ and MT where observed for increasing ergosterol content in the membrane (37). However it should be noted that, at a significance level of *p* = 0.01, there are no statistically significant differences in κ between the populations containing 0% and 10% ergosterol, nor between those containing 30% and 50% ergosterol. Both composition, with 0% and 10% ergosterol are in agreement with κ established for Yeast Extract membrane (Fig 3B). However, presence of ergosterol did influence other investigated parameters, hence inclusion of small ergosterol fraction might contribute to better agreement of the model. Finally, a reduction in lateral diffusion was detected, consistent with the known capacity of cholesterol to diminish overall membrane fluidity (39).

**Figure 3:**
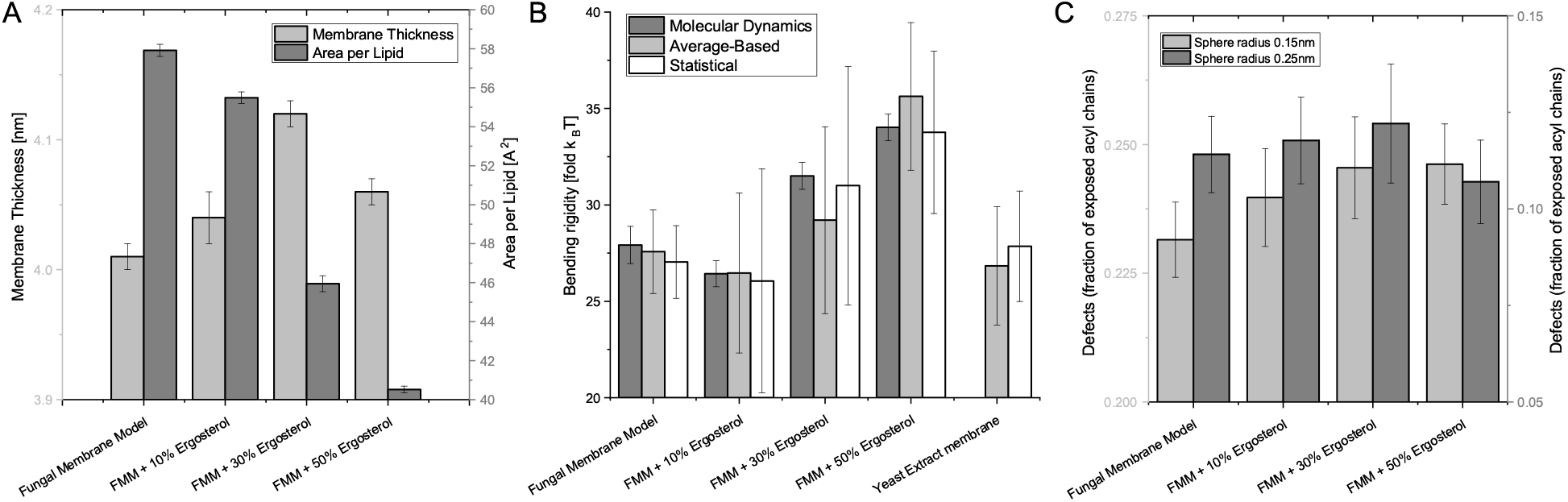
Biophysical characteristics of the investigated fungal plasma membrane model (FPMM) with addition of ergosterol. (A) Membrane thickness (MT) and area per lipid (APL) determined from molecular dynamics (MD) simulations. (B) Bending rigidity (κ) determined from MD simulations and experimentally via flicker-noise spectroscopy using two different calculation approaches. (C) Acyl chain accessibility sphere analysis used to quantify membrane packing defects for the investigated systems at two probe radii.

The FTIR spectra of membranes containing 50% ergosterol suggest that it has a limited influence on the overall spectral profile, indicating that its incorporation into the lipid membrane may be less effective than expected. In the 700 − 1800 cm^−1^ region, only minor spectral changes are observed relative to the five-component membrane, while the characteristic phospholipid bands remain largely unchanged. See Fig S4 for detailed spectra. The most pronounced differences are detected in the 810 − 1260 cm^−1^ region, where we can detect ERG for PC:PE:PI:PA:PS+10%ERG membrane. The characteristic band observed near 2600 cm^−1^ is clearly detectable in the membrane containing 10% ERG; however, it is absent in the 50% ERG formulation. This may indicate that band is substantially weakened or masked by overlapping. It might also suggest that increasing the ergosterol content does not produce proportional spectral changes or strongest rearangment of the membrane may occurs in 10-30% ERG concentration. However, this result strengthens the message of including a small ergosterol fraction in the membrane model, as it does affect the membrane spectra in various bands.

The impact of ergosterol on the lipid membrane defects was subtle. In general, the addition of ergosterol increases acyl chain accessibility, which contrasts with the typical condensing and ordering behavior observed for sterols(40). However, in the FPMM system containing 50 mol% ergosterol, the frequency of small packing defects (*r* = 0.25 nm) decreased, differing from the behavior observed at lower concentrations (10 mol% and 30 mol%; Fig 3C). This observation supports the ATR-FTIR analysis, which revealed the absence of the 2600 cm^−1^ band in the 50 mol% ergosterol composition. At lower concentrations, these changes were marginal; statistical testing (α = 0.01) indicated no significant differences between the control FPMM and FPMM with 10 mol% ergosterol, nor between the 10 mol% and 30 mol% compositions. Overall, low-to-moderate ergosterol levels do not substantially alter defect formation in FPMM. This is likely because the pre-existing PE–PI/PA hydrogen-bonding networks already establish lateral domain segregation, thereby inhibiting the typical rigidifying effect of ergosterol on membrane packing.

The non-monotonic impact of ergosterol on FPMM observed in FTIR and MD data is qualitatively consistent with non-monotonic behavior reported with neutron diffraction and neutron spin echo techniques (38), although the exact MT and κ trends have not been replicated. However, this might be due to the more complex interaction of ergosterol with FPMM, characterized by an intricate environment of various lipid species, compared to the simple ergosterol/POPC model used in (38).

### 2.3. Effect of Triglycerides

Triacylglycerols function as metabolic buffers in the fungal plasma membrane, regulating its fluidity and preventing structural collapse under environmental stress. They safely store excess, toxic free fatty acids that would otherwise dissolve the membrane bilayer. In addition, they serve as an immediate, on-demand reservoir of fatty acids required for rapid membrane remodeling and hyphal growth. The *Histoplasma* and *Penicillium* membranes consisted of 45% and 30% triacylglycerol, respectively (41; 42).

TG has a significant effect on membrane thickness, most likely due to a combination of acyl chain straightening and decrease of inter-leaflet coupling. Indeed, in simulated system interdigitation decreased significantly, from 0.58 ± 0.02 nm for the control membrane (0% TG) down to 0.32 ± 0.01 nm for the 45% TG membrane. Interestingly, the effect on MT is most pronounced at 15% and 30% TG, whereas the change between 30% and 45% TG is relatively small. TG incorporation also led to a decrease in APL, which is surprising given the bulky size of this molecule. Indeed, as can be seen in Fig 4B, the TGs localize mostly in the acyl chain region, also significantly altering contact between other lipids acyl chains, as indicated by changing interdigitation. These structural alterations provide a compelling biophysical rationale for the role of TGs as metabolic and structural buffers in fungal plasma membranes. The observed increase in MT combined with reduced APL reinforces the hydrophobic core, enhancing resistance against osmotic and mechanical stresses that could otherwise cause structural collapse. Simultaneously, the pronounced drop in interdigitation decouples the opposing leaflets, allowing individual leaflets to slip laterally and thereby maintaining functional membrane fluidity despite tighter acyl-chain packing.

**Figure 4:**
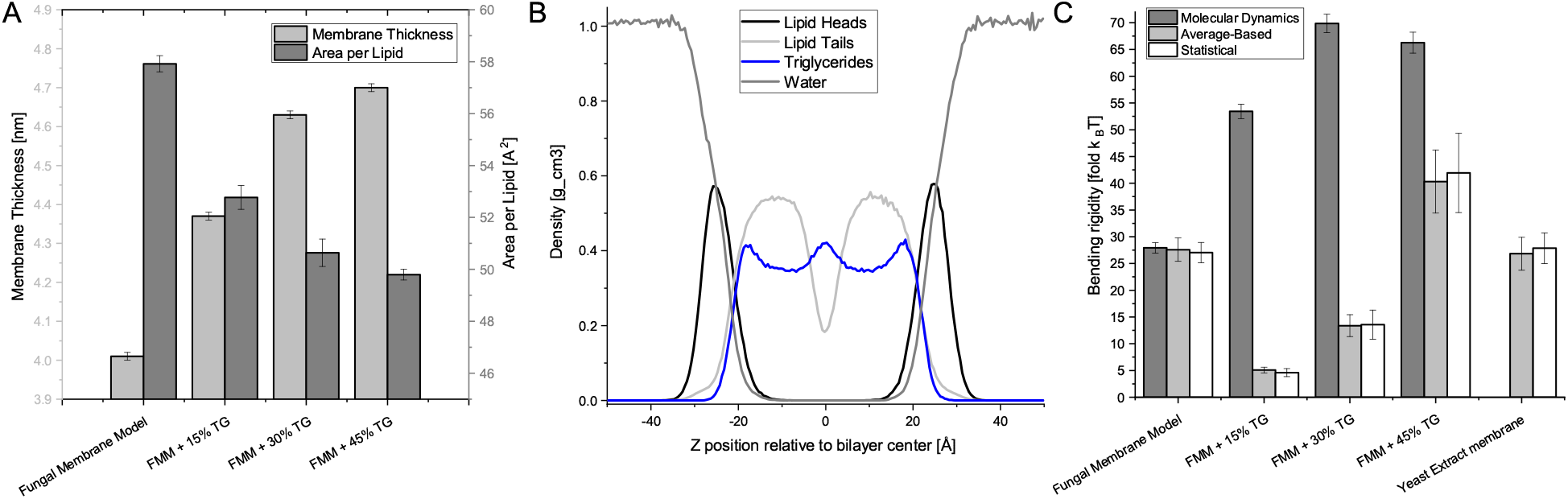
Biophysical characteristics of the investigated fungal plasma membrane model (FPMM) with addition of triacylglycerols. (A) Membrane thickness (MT) and area per lipid (APL) determined from molecular dynamics (MD) simulations. (B) Mass density profile across FPMM + %30TG membrane normal. (C) Bending rigidity (κ) determined from MD simulations and experimentally via flicker-noise spectroscopy using two different calculation approaches.

When analyzing the impact of TGs on membrane mechanics – specifically bending rigidity – a significant discrepancy occurred between the *in silico* and *in vitro* approaches. In the MD simulations, a substantial increase in bending rigidity (approximately 2-fold) was observed following the addition of 15% TG. At 30% TG, the bending rigidity increased further, though less dramatically (from 53.4 ± 1.4 *k*_B_*T* to 69.9 ± 1.7 *k*_B_*T*), whereas at 45% TG, it slightly decreased to 66.0 ± 2.0 *k*_B_*T* . On the other hand, the experimental measurements revealed an entirely opposite trend. Upon the addition of 15% TG, the bending rigidity decreased significantly to 4.6 ± 0.8 *k*_B_*T* compared to the experimental baseline for FPMM. In the 30% TG system, the value was slightly higher (13.5 ± 2.71 *k*_B_*T*), before increasing sharply at 45% TG to reach 41.9 ± 7.4 *k*_B_*T* . This latter value was nearly 2-fold higher than the baseline FPMM system and 3-fold higher than the 30% TG system. Similar discrepancies between *in silico* and *in vitro* observations previously have been reported (43). Most likely explanation of system not reaching equilibrium has been rejected by running simulated system with 30% TG for additional 200 ns – no change in bending was observed. A plausible explanation of this discrepancy might be due to the molecular dimensions of TGs. Within MD simulations, periodic boundary conditions limits lipid rearrangements and splay fluctuations within a finite unit cell. The presence of bulky triacylglycerols can generate local packing stresses, but the periodic membrane geometry restricts how these orientational perturbations can propagate and relax beyond the simulation box. As adapted real-space fluctuation method determines bending rigidity from lipid splay fluctuations, this can decrease the measured fluctuation amplitude and result in an overestimated bending modulus. Indeed, the effect of TG on the membrane was reported to be closer to the experimental trend. The significant decrease in membrane bending rigidity upon addition of small fraction of TGs has been reported in other study employing vesicle fluctuation spectroscopy, where POPC membrane enriched with 10% of triolein exhibited almost 3-fold lower κ compared to reference, pure POPC membrane (44). Similar findings have been reported in nanotube pulling experiments, were the presence of triolein decreased membrane bending rigidity by a factor of two (45).

ATR-FTIR analysis confirmed the presence of TG in the system. Specifically, the intensity of the absorption bands in the 700 − 1800 cm^−1^ region increased with increasing TG concentration. In contrast, the intensity of the lipid CH stretching region 3050 − 2800 cm^−1^ decreased, reflecting the reduced relative contribution of phospholipid vibrations to the overall spectrum as the TG content increased. The spectral shifts observed in the 1730 cm^−1^ region agree well with the trends measured for *K*_*A*_. However, changes within the 1670 cm^−1^ region do not directly correspond to the experimental bending rigidity values, suggesting instead a monotonic increase in bending stiffness. This discrepancy for the 1673 cm^−1^ band may result from the reduced spatial freedom of the lipid film compared to a fully hydrated, aqueous environment. Because the lipid films analyzed by ATR-FTIR undergo dehydration during preparation, interfacial lipid mobility and long-range conformational fluctuations become physically constrained. Consequently, the mechanical state of the dehydrated film aligns more closely with that of the simulated MD system – where spatial fluctuations are similarly restricted by periodic boundary conditions – resulting in a consistent increase in bending rigidity.

Finally, in each of the investigated TG concentrations, lateral domains were clearly present. FTIR results show that although the intensity of the broad 3500–3050 cm^−1^ band (assigned to O–H and N–H stretching vibrations) was lower than that observed for the five-component control membrane without TG, it remained relatively high. This suggest that hydrogen-bonding interactions were largely preserved despite TG incorporation, thereby supporting the sustained presence of these domains. The TG presence in membrane also had power trend effect on packing defects. For 15% TG no effect was observed, but for 45% the increase of packing defects was significant (see Fig S5).

## 3. Materials and methods

### 3.1. Materials

Lipids: POPC (1-palmitoyl-2-oleoyl-glycero-3-phosphocholine), DYPE (1,2-di-(9Z-hexadecenoyl)-snglycero-3-phosphoethanolamine), DSPS (1,2-dioctadecanoyl-sn-glycero-3-phospho-L-serine), DPPI (1,2-dihexadecanoyl-sn-glycero-3-phospho-(1’-myo-inositol)), DLiPA (1,2-di-(9Z,12Z-octadecadienoyl)-sn-glycero-3-phosphate), PPST (1,3(d5)-dihexadecanoyl-2-octadecanoyl-glycerol), Yeast Polar Extract and Ergosterol were purchased from Avanti Polar Lipids (Alabaster, AL, USA). Fluorescent probes: TexasRed DHPE was purchased from ThermoFisher (International) and Atto488-DOPE was purchased from Atto-Tech (Siegen, Germany).

### 3.2. Molecular Dynamics Simulations and Parametrization

The full-atomistic MD simulation was performed using NAMD 2.13 (46) software with CHARMM36 force fields (47) under NPT conditions (constant: Number of particles, Pressure and Temperature). Lipid membrane systems consisted of 648 lipid molecules (324 on each of leaflets). Following systems were simulated – POPC:DYPE (60:40), POPC:DYPE:DSPS (52:33:15), POPC:DYPE:DPPI (56:37:7), POPC:DYPE:DLiPA (55:36:9), POPC:DYPE:DPPI:DSPS (51:33:7:9), POPC:DYPE:DSPS:DLiPA (47:31:8:14), POPC:DYPE:DPPI:DLiPA:DSPS (44:29:13:8:6). Furthermore, quinary mixture model was also simulated with ergosterol (10,30 and 50%) as well as with triglyceride (15,30,45%). This resulted in total 14 system (details are presented in Fig S6 and Table S3). Each system was hydrated in such a way that 75 water molecules per lipid molecule was used. All systems were additionally neutralized with *Na*^+^ ions. All systems were equilibrated using CHARMM equilibration procedure (48). Total simulation time for all systems was at least 110 ns with last 10 ns used for analysis, each system had 3 replicas performed - presented values are average of those three replicas. Our previous results indicate such simulation times should be sufficient for quinary system to accurately reflect biophysical properties (49). Simulations were carried under 25°C (298.15K) to maintain agreement with experimental measurements.

### 3.3. Membrane system characteristics

All biophysical parameters were calculated in accordance with the methodologies detailed in our previous work (49). Briefly, area per lipid (APL) and membrane thickness (MT) were determined via Voronoi tessellation and leaflet-to-leaflet atomic plane distances using custom MATLAB scripts. Bending rigidity (*κ*) and tilt modulus were evaluated via the real-space fluctuation (RSF) method (50). Area compressibility (*K*_*A*_) was derived from local thickness fluctuations following the framework by Doktorova *et al*. (51). Lateral diffusion coefficients (*D*_2*D*_) were obtained from mean square displacements (MSD) via the Diffusion Coefficient Tool (52), and inter-leaflet mass overlap (interdigitation) was calculated using MEMB-PLUGIN (53). A complete description of each calculation protocol is available (49). Finally, membrane packing defects (acyl chain accessibility) were evaluated following the protocol by Boyd *et al*. (29), employing a grid-based surface occupancy map (0.5 Å spacing) combined with sphere-probe analysis (radii 0.15 and 0.25 nm) to identify hydrophobic surface patches. The complete procedural details and the atom selection criteria are provided in (30). To quantify the local lipid composition surrounding PA lipids, spatial proximity analysis was performed across the MD trajectory using a custom Python script built on the MDAnalysis framework (v2.9.0) (54). For each frame, interatomic distances between the P atoms of PA and neighboring lipid species were calculated. A spatial cutoff criterion of 12.0 Å (measured between phosphorus headgroup coordinates) was used to define immediate lipid–lipid contacts. Hydrogen-bond analysis between lipid molecules was performed using the HydrogenBondAnalysis module implemented in MDAnalysis. Hydrogen bonds were identified based on a donor–acceptor distance cutoff of 3.0 Å and a donor– hydrogen–acceptor angle cutoff of 150^°^. Only intermolecular hydrogen bonds between distinct lipid residues were considered, while intramolecular interactions were excluded.

### 3.4. GUVs Electroformation

The modified method of model membrane formation for giant unilamellar vesicles (GUV) was used. Briefly, 1 mM lipid solution with fluorescent probe (1m%) in chloroform was distributed equally along the platinum electrodes with density of 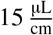 and dried under reduced pressure for 2 hours. The electrodes were then submerged in 100 mM sucrose solution and a square 1 Hz AC electric field was applied for 23 h. The protocol of electroformation involved increasing the voltage from 2 V by 2 V every hour until reaching 8 V, which has been applied for the remaining 20 h. For FTMM with either ergosterol or triglycerides the protocol has been modified – as it yielded better results – with total electroformation lasting 2.5 h, which involved applying a 500 Hz square AC electric field with a series of increasing voltage steps: 0.14 V, 1.25 V, 5 V, lasting respectively 600 s, 1200 s and 5400 s. For the remaining 30 min the frequency has been reduced to 1 Hz. The electroformation has been conducted in temperature exceeding *T*_m_ of all lipids in the mixture, which involved usage of custom heated glass chambers for temperatures over 45 ^°^C. It should be noted that the elevated temperature during electroformation proved unnecessary for the investigated mixtures; the DPPI content in the final vesicles remained identical regardless of whether heating was applied (Fig S7). For lower or equal temperatures custom PTFE (polytetrafluoroethylene) chambers were used.

### 3.5. Flicker Noise spectroscopy

Thermally induced shape fluctuations of GUVs were used to determine mechanical properties (bending rigidity and surface tension). The series of images of GUVs was recorded by camera-based fluorescence microscopy. A Leica TCS SPE (Leica, Wetzlar, Germany) was equipped with a 63×/1.30 ACS APO oil-immersion objective. Fluorophores were excited with an EL6000 illuminator (mercury metal halide bulb), and fluorescence emission was detected using I3 cut-off filter (BP 450-490 nm excitation filter, a 510 nm dichromatic mirror, and an LP 515 nm emission filter, transmitting light above 515 nm). Time-series movies were acquired at 1392×1040 pixels using a Leica DFC310 FX camera, typically comprising 3000 consecutive frames, with the imaging plane positioned at the vesicle center to maximize contour fidelity. The two-dimensional liposome images are transformed to the three-dimensional Helfrich model using both the average-based and statistical approaches. Then the radial position of the bilayer, extracted from images, is used to construct angular autocorrelation curves. In the average-based approach autocorrelation curves are decomposed into Legendre polynomial series and are plotted as a function of fluctuation mode so the bending rigidity coefficient can be determined. In the statistical approach autocorrelation curves are decomposed into Fourier series and a frequency histogram of amplitudes for each mode of fluctuation is calculated. The histogram is then used for determination of the bending rigidity coefficient. The radii of investigated vesicles ranged from 2.9 up to 19.1 *µm*.

### 3.6. ATR-FTIR spectroscopy

ATR-FTIR spectra were recorded using a Nicolet Magna-860 FTIR spectrometer equipped with a Golden Gate single-reflection heated diamond ATR accessory (Specac Inc.). Measurements were performed on thin lipid films. Briefly, GUVs electroformed in water were placed overnight at 95^°^C to evaporate the aqueous solvent. Each dry sample was subsequently resuspended in 100 *µ*L of ultrapure water and transfered on diamond surface. Lipid films were formed by a slow evaporation of water under a stream of air. Spectra were collected in the 4000 − 400 cm^−1^ region by accumulating 128 scans at a spectral resolution of 1 cm^−1^. All measurements were conducted at room temperature. Reference spectra are available in Fig S8. Raw spectra were pre-processed to ensure reproducibility across measurements. First, spectral smoothing was applied using a Savitzky-Golay filter with a 19-point window. Subsequent baseline correction was performed using seven fixed anchor points to maintain a consistent baseline across all spectra. Following baseline correction, the spectra were normalized to the CH_2_ band by dividing the entire spectrum by the integrated area of this band (calculated between 1450 and 1480 cm^−1^ relative to zero absorbance). Band area integration, rather than peak maximum height, was selected as the normalization reference because integration over the full band envelope minimizes sensitivity to single-point random noise and minor peak shifts, which could otherwise compromise single-point maximum readings(55). Data analysis and spectral processing were performed using OriginPro 2024 (OriginLabs).

### 3.7. Statistics

In order to test for the significant difference between the parameters, unless specified otherwise, the one-way ANOVA test was used with the significance level at 0.05. The Tukey test was used as a post hoc test. All statistical analysis was performed using the OriginPro 2024 (OriginLabs). Average values are presented with standard deviation.

## 4. Summary and conclusions

In this study, a new fungal plasma membrane model (FPMM) was proposed and evaluated. Using a bottom-up approach, a detailed biophysical characterization of each constituent lipid mixture was performed by combining *in silico* (molecular dynamics) and *in vitro* techniques (flicker noise spectroscopy and ATR-FTIR spectroscopy). Findings from these complementary approaches demonstrate that the proposed POPC:DYPE:DSPS:DLiPA:DPPI quinary lipid mixture accurately models key biophysical characteristics of the fungal plasma membrane.

Among all investigated lipids, the independent presence of DPPI – as well as its specific interactions with DSPS – induced the most substantial deviations in baseline membrane properties, particularly regarding APL, MT, and *K*_*A*_. On the other hand, DSPS played a dominant role in modulating membrane bending rigidity (*κ*). The highly unsaturated DLiPA did not significantly alter baseline physical parameters, though it exerted the most pronounced impact on membrane packing defects. Notably, the overall biophysical profile of the proposed FPMM closely mimicked that of natural vesicles derived from yeast lipid extracts.

Additionally, the roles of ergosterol and triacylglycerols (TGs) on proposed FPMM were characterized. The addition of ergosterol induced notable changes in membrane properties, increasing both mechanical properties (*K*_*A*_ and *κ*), although its impact on bending rigidity at lower concentrations and on packing defect frequency remained subtle. Nevertheless, because ergosterol influences interfacial organization, its inclusion at physiological fractions enhances the mimetic fidelity of the FPMM model toward native fungal membranes. In contrast, TGs caused marked alterations across all biophysical parameters. A discrepancy between experimental and computational bending rigidity values was identified in TG-containing systems, highlighting the inherent limitations of periodic boundary conditions in capturing fluctuation modes in the presence of bulky hydrophobic molecules.

## Acknowledgements

This work was possible thanks to the financial support from the National Science Centre (Poland) grant 2024/55/D/NZ9/01631.

