## Supplementary Information Figures and Tables for "A Bottom-Up Approach to Fungal Plasma Membrane Model: Lipid Mixture Design and Biophysical-Mechanical Characterization"

---

---

### 1. Model building

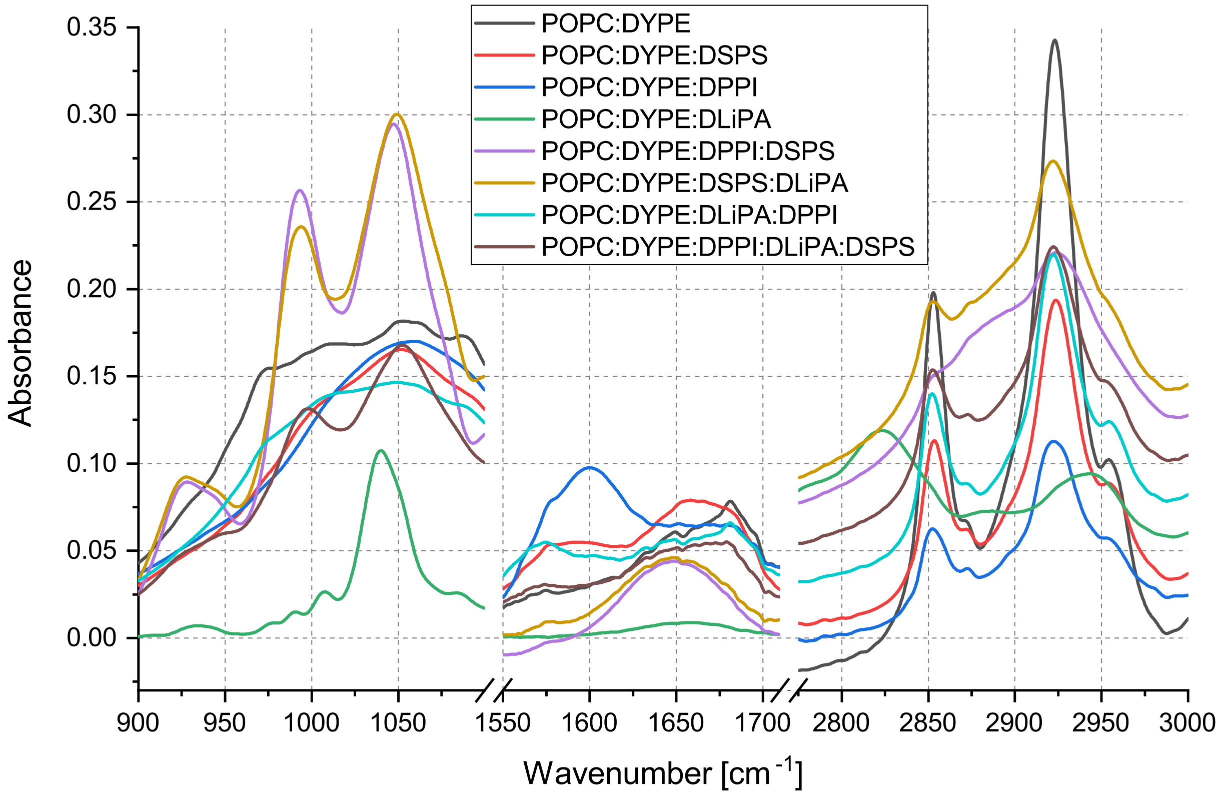


Figure S1: Absorbance spectra obtained from ATR-FTIR spectroscopy.Selected three most relevant bands are presented.

Table 1: Membrane parameters calculated from Molecular Dynamics simulation. Each value is average and standard deviation from three replicas.

| Membrane Mixture and Molar Ratio | Membrane thickness [nm] | Area per lipid [Å$^2$] | bending rigidity [$J \cdot 10^{-19}$] | tilt rigidity [$J \cdot 10^{-20}$] | $K_A$ [mN/m] | Lateral diffusion [$\mu m^2/s$] | Intdgt [nm] |
|---|---|---|---|---|---|---|---|
| PC:PE 60:40 | 3.88 ± 0.01 | 59.57 ± 0.19 | 1.18 ± 0.02 | 3.28 ± 0.11 | 359 ± 53 | 5.4 ± 0.2 | 0.60 ± 0.01 |
| PC:PE:PS 52:33:15 | 3.87 ± 0.01 | 60.34 ± 0.33 | 1.06 ± 0.03 | 3.32 ± 0.12 | 360 ± 35 | 5.2 ± 0.4 | 0.60 ± 0.02 |
| PC:PE:PI 56:37:7 | 3.94 ± 0.02 | 58.56 ± 0.67 | 1.15 ± 0.03 | 3.67 ± 0.07 | 498 ± 49 | 4.5 ± 0.4 | 0.52 ± 0.02 |
| PC:PE:PA 55:36:9 | 3.88 ± 0.01 | 59.79 ± 0.12 | 1.13 ± 0.05 | 3.25 ± 0.15 | 311 ± 49 | 5.52 ± 0.34 | 0.55 ± 0.02 |
| PC:PE:PI:PS 48:32:14:6 | 3.98 ± 0.01 | 58.16 ± 0.37 | 1.15 ± 0.04 | 3.63 ± 0.08 | 491 ± 35 | 4.15 ± 0.13 | 0.51 ± 0.00 |
| PC:PE:PS:PA 51:33:7:9 | 3.93 ± 0.01 | 59.07 ± 0.47 | 1.07 ± 0.01 | 3.31 ± 0.07 | 403 ± 73 | 4.8 ± 0.2 | 0.57 ± 0.02 |
| PC:PE:PA:PI 47:31:8:14 | 3.94 ± 0.02 | 58.56 ± 0.80 | 1.12 ± 0.03 | 3.56 ± 0.10 | 462 ± 59 | 4.5 ± 0.1 | 0.54 ± 0.05 |
| PC:PE:PI:PA:PS (FPMM) 44:29:13:8:6 | 4.01 ± 0.01 | 57.92 ± 0.31 | 1.14 ± 0.03 | 3.51 ± 0.03 | 445 ± 99 | 4.4 ± 0.3 | 0.58 ± 0.02 |

Abbreviations: PC stands for POPC; PE - DYPE; PS - DSPS; PI - DPPI; PA - DLiPA; FPMM - fungal plasma membrane model; $K_A$ - area compressibility; Intdgt - interdigitation;

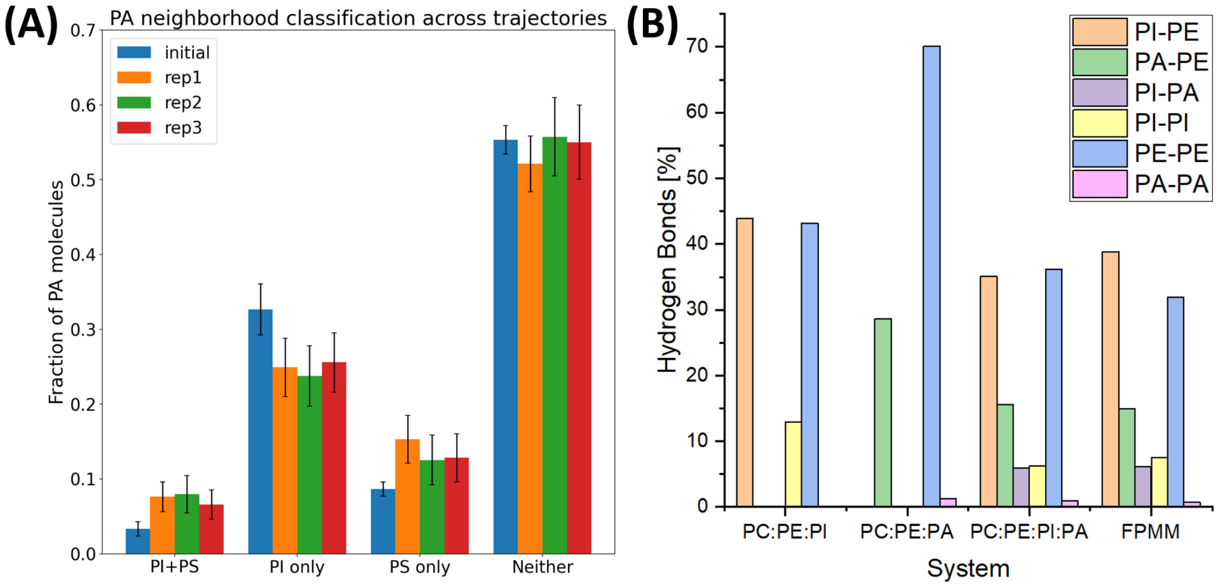


Figure S2: Graph description

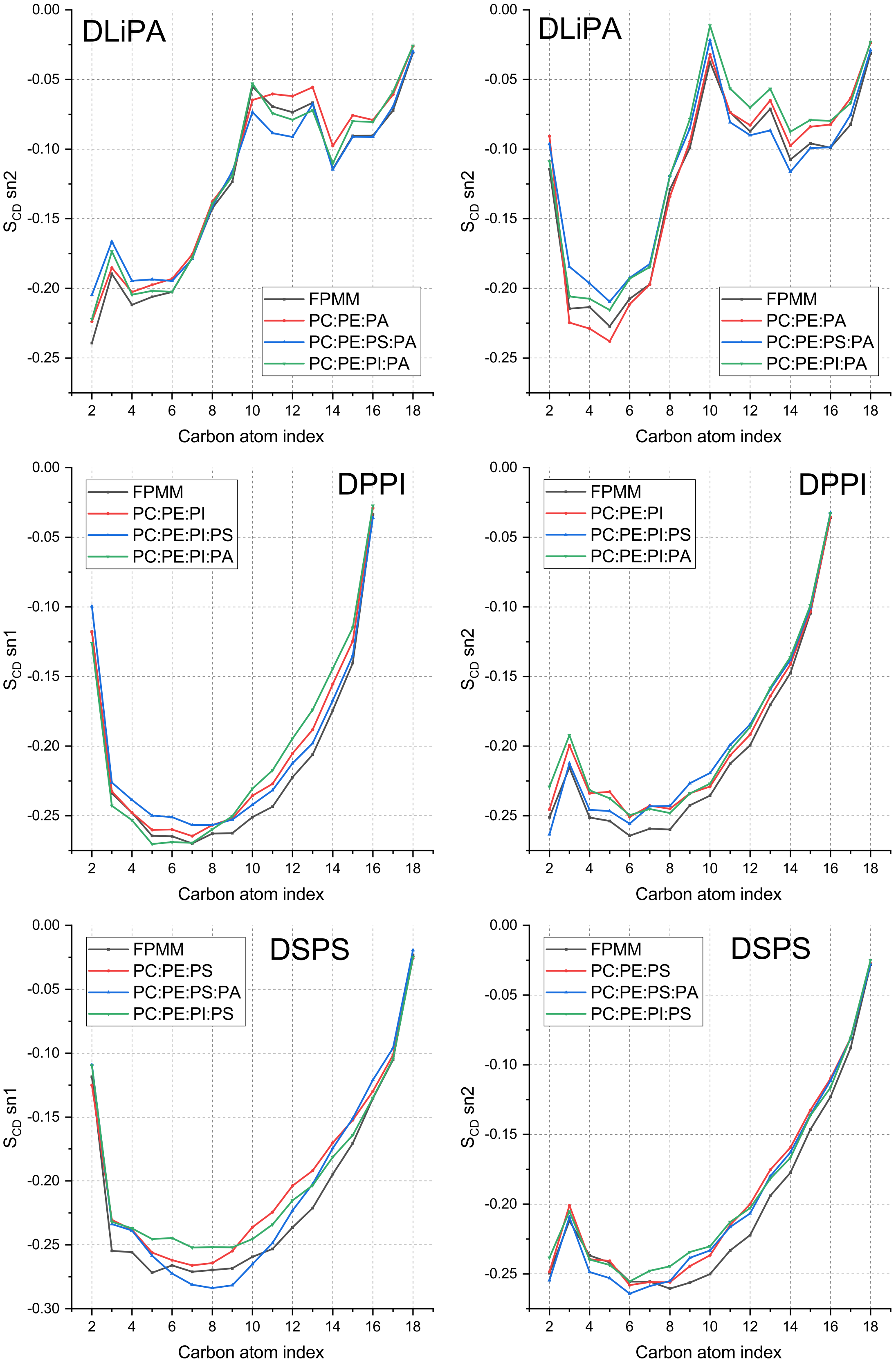




Figure S3: Order parameter ($S_{CD}$ values) for lipids DLiPA, DPPI and DSPS in different membrane systems.

### 2. Ergosterol and TGs

Table 2: Membrane parameters for different ergosterol and triglyceride concentrations calculated from Molecular Dynamics simulation. Each value is average and standard deviation from three replicas.

| Membrane Mixture | Membrane Thickness [nm] | Area per lipid [Å$^2$] | bending rigidity [$J \cdot 10^{-19}$] | tilt rigidity [$J \cdot 10^{-20}$] | $K_A$ [mN/m] | Lateral diffusion [$\mu m^2/s$] | Intdgt [nm] |
|---|---|---|---|---|---|---|---|
| FPMM | 4.01 ± 0.01 | 57.92 ± 0.31 | 1.14 ± 0.03 | 3.51 ± 0.03 | 445 ± 99 | 4.4 ± 0.3 | 0.58 ± 0.02 |
| FPMM + 10% ERG | 4.04 ± 0.02 | 55.49 ± 0.30 | 1.08 ± 0.01 | 3.89 ± 0.20 | 543 ± 262 | 4.1 ± 0.2 | 0.48 ± 0.01 |
| FPMM + 30% ERG | 4.12 ± 0.01 | 45.95 ± 0.41 | 1.28 ± 0.04 | 5.24 ± 0.05 | 920 ± 209 | 3.2 ± 0.2 | 0.40 ± 0.01 |
| FPMM + 50% ERG | 4.06 ± 0.01 | 40.52 ± 0.17 | 1.389 ± 0.06 | 6.13 ± 0.19 | 1996 ± 640 | 3.1 ± 0.2 | 0.34 ± 0.01 |
| FPMM + 15% PPSTG | 4.37 ± 0.01 | 52.77 ± 0.46 | 2.43 ± 0.12 | 7.88 ± 0.64 | 799 ± 630 | 2.6 ± 0.2 | 0.43 ± 0.01 |
| FPMM + 30% PPSTG | 4.63 ± 0.01 | 50.64 ± 0.53 | 4.73 ± 0.19 | 20.14 ± 0.56 | 1849 ± 374 | 2.2 ± 0.2 | 0.39 ± 0.02 |
| FPMM + 45% PPSTG | 4.70 ± 0.01 | 49.80 ± 0.21 | 5.31 ± 0.18 | 32.46 ± 0.29 | 3832 ± 732 | 1.3 ± 0.1 | 0.32 ± 0.01 |

Abbreviations: FPMM stands for fungal plasma membrane model; ERG - ergosterol; PPSTG - palmitic acid (16:0), palmitic acid (16:0), stearic acid (18:0) triglyceride; $K_A$ - area compressibility; Intdgt - interdigitation;

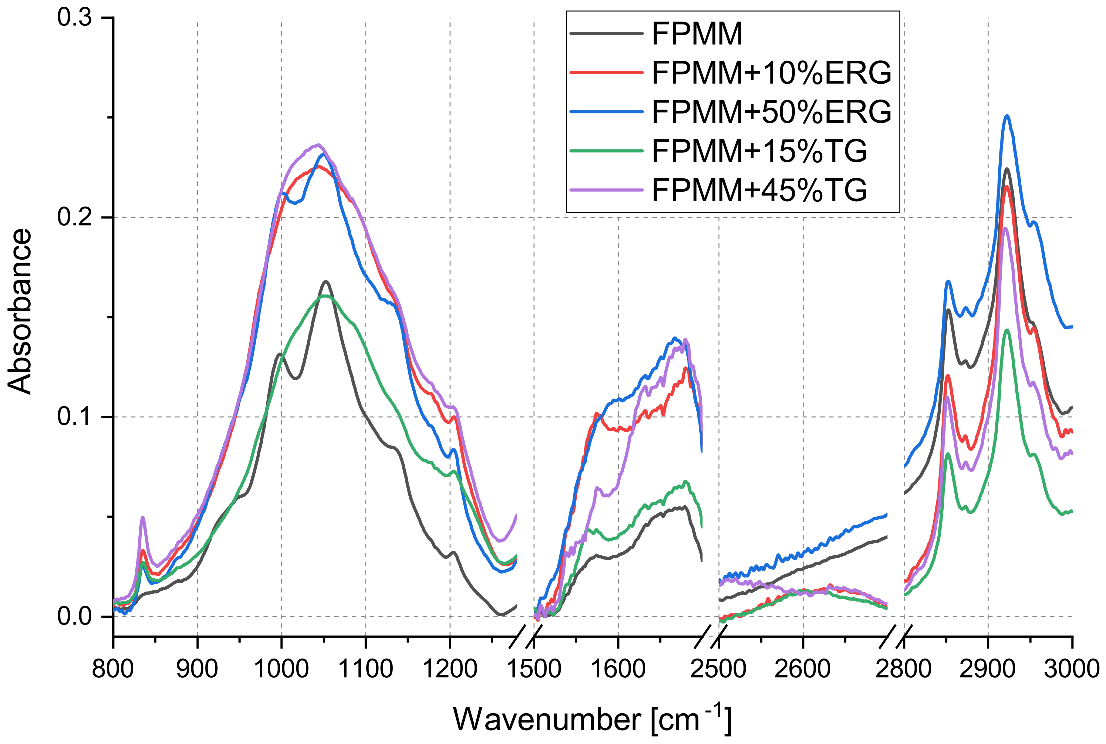


Figure S4: Absorbance spectra obtained from ATR-FTIR spectroscopy for FPMM (fungal plasma membrane model) and different concentration of Ergosterol and/or TGs.Selected three most relevant bands are presented.

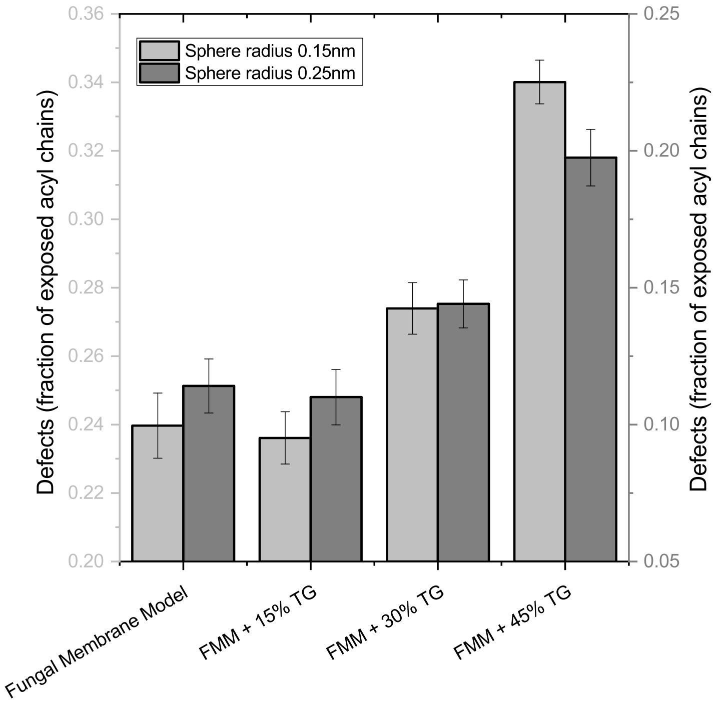


Figure S5: Acyl chain accessibility sphere analysis used to quantify membrane packing defects for the investigated systems at two probe radii for FPMM system with different addition of TGs.

### 3. Molecular dynamics system setup

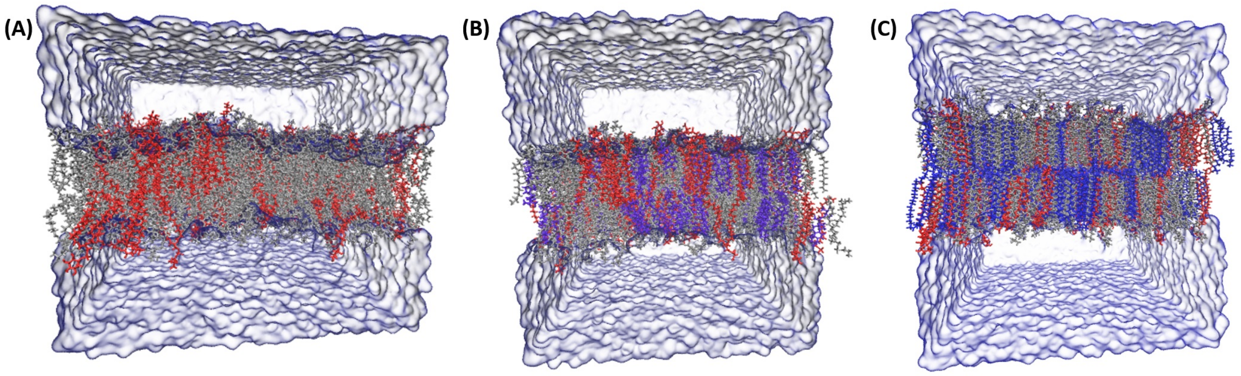


Figure S6: Snapshots of (A) FPMM, (B) FPMM + 30% Ergosterol and (C) FPMM + 30% PPSTG systems after molecular dynamics simulation. The coloring is as follows: POPC and DYPE lipids are colored silver, DLiPA DPPI and DSPS lipids are colored red, ergosterol molecules are colored violet and PPSTG are colored blue.

Table 3: DESC

| System | Number of molecules | | | | | | | |
|---|---|---|---|---|---|---|---|---|
| | PC | PE | PS | PI | PA | ERG | PPSTG | Ions $Na^{+}$ |
| POPC:DYPE | 388 | 260 | 0 | 0 | 0 | 0 | 0 | 0 |
| POPC:DYPE:DSPS | 334 | 218 | 96 | 0 | 0 | 0 | 0 | 96 |
| POPC:DYPE:DPPI | 364 | 238 | 0 | 46 | 0 | 0 | 0 | 0 |
| POPC:DYPE:DLiPA | 356 | 232 | 0 | 0 | 60 | 0 | 0 | 60 |
| POPC:DYPE:DPPI:DSPS | 314 | 204 | 40 | 90 | 0 | 0 | 0 | 130 |
| POPC:DYPE:DSPS:DLiPA | 334 | 216 | 42 | 0 | 56 | 0 | 0 | 98 |
| POPC:DYPE:DLiPA:DPPI | 308 | 200 | 0 | 88 | 52 | 0 | 0 | 140 |
| FPMM | 290 | 190 | 36 | 82 | 50 | 0 | 0 | 168 |
| FPMM:ERG10 | 262 | 170 | 34 | 74 | 44 | 64 | 0 | 152 |
| FPMM:ERG30 | 204 | 132 | 26 | 58 | 34 | 194 | 0 | 118 |
| FPMM:ERG50 | 146 | 94 | 18 | 42 | 24 | 324 | 0 | 84 |
| FPMM:PPSTG15 | 246 | 160 | 32 | 70 | 42 | 0 | 98 | 144 |
| FPMM:PPSTG30 | 202 | 132 | 26 | 58 | 34 | 0 | 196 | 118 |
| FPMM:PPSTG45 | 158 | 104 | 20 | 46 | 28 | 0 | 292 | 94 |

### 4. Electroformation Heating

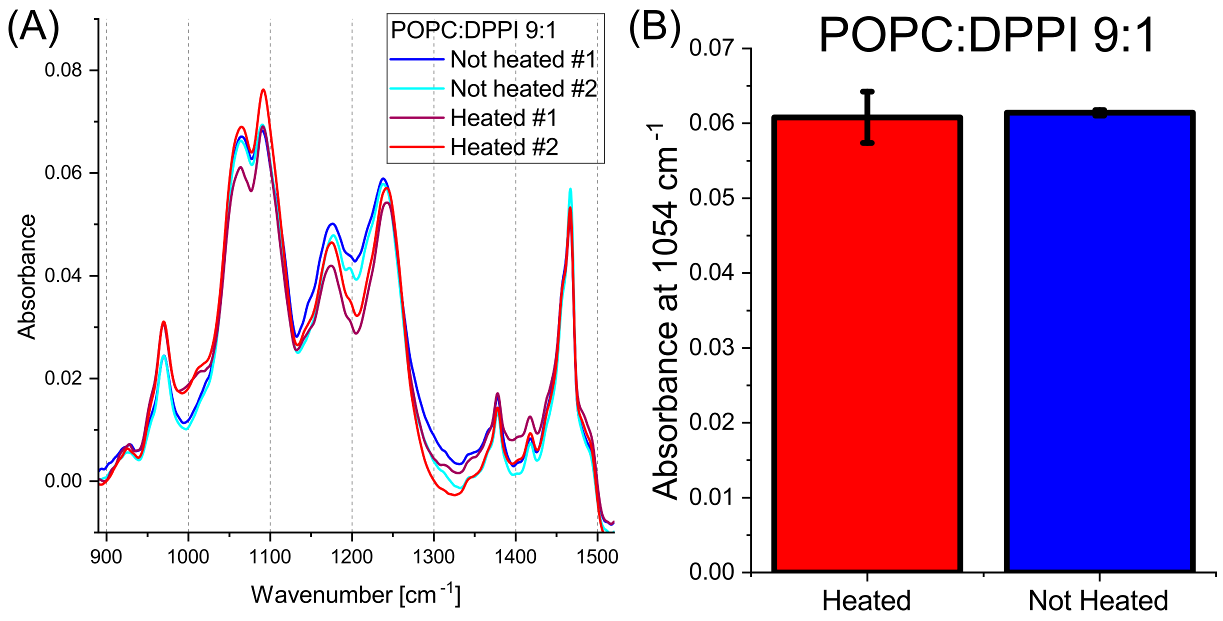


Figure S7: (A) Absorbance spectra obtained from ATR-FTIR spectroscopy for two distinct ways of electoforming POPC:DPPI 9:1 GUVs - with heating and without heating. (B) Absorbance at 1054$cm^{-1}$, which is a band strongly contributioned from the inositol ring of DDPI. Lack of significant change indicates no difference in DPPI concentrations in GUVs regardless of heating state.

### 5. ATR-FTIR single component references

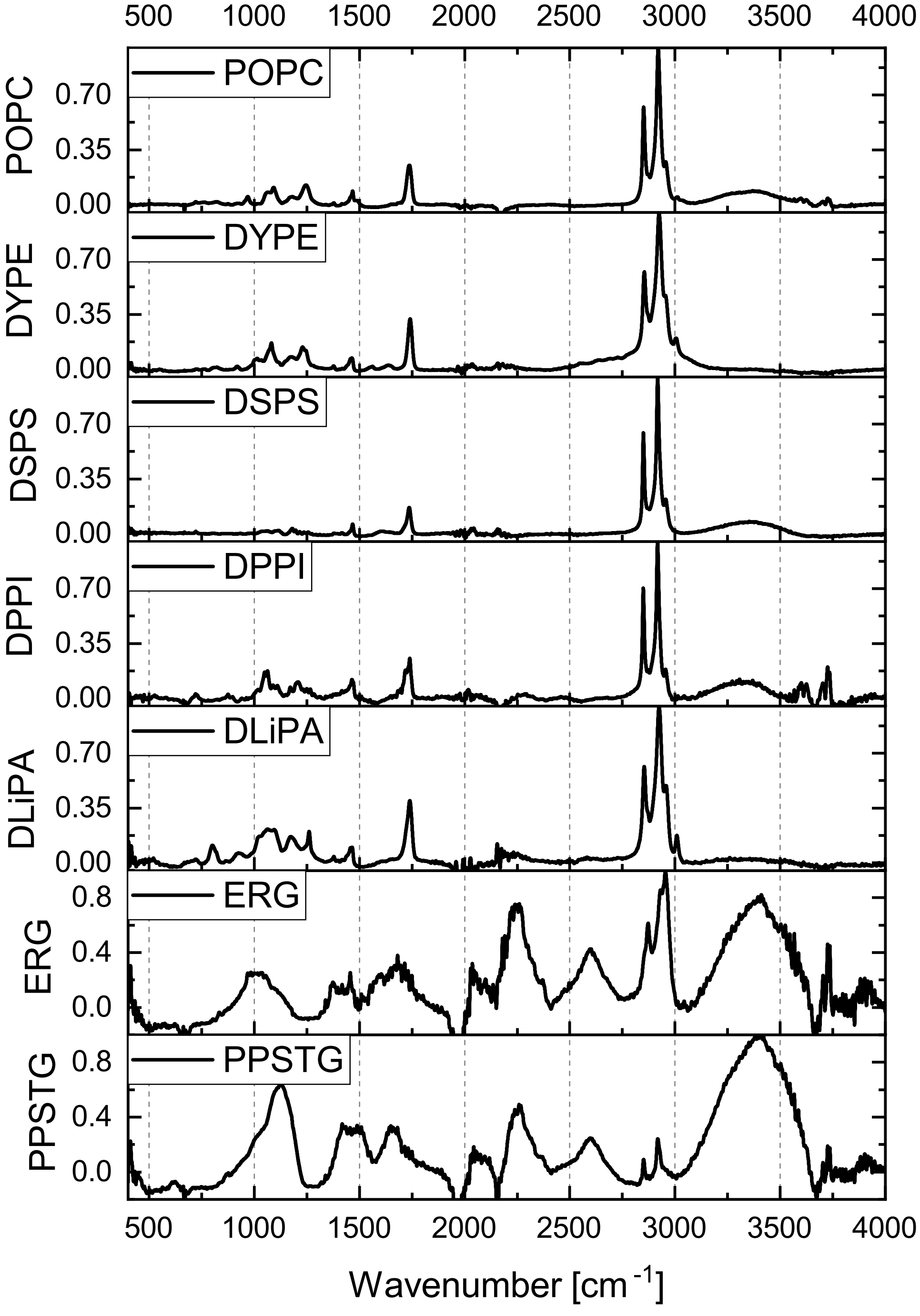


Figure S8: Normalized ATR-FTIR reference spectra of individual lipid components constituting the membrane, as well as investigated ergosterol and triglyceride.
